## Extended Figures for "Chemotherapeutic 6-thio-2’-deoxyguanosine selectively targets and inhibits telomerase by inducing a non-productive telomere-bound telomerase complex"

### Extended Figure 1

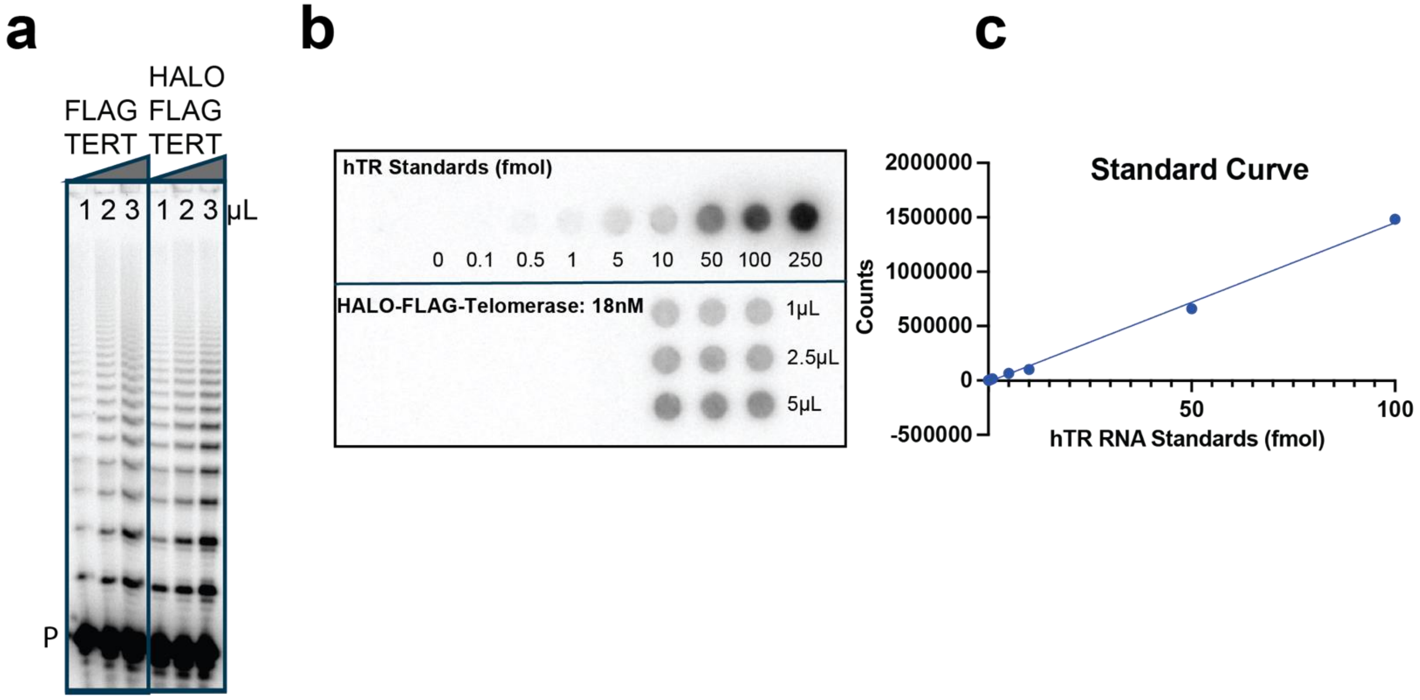

### Extended Figure 2

**a**

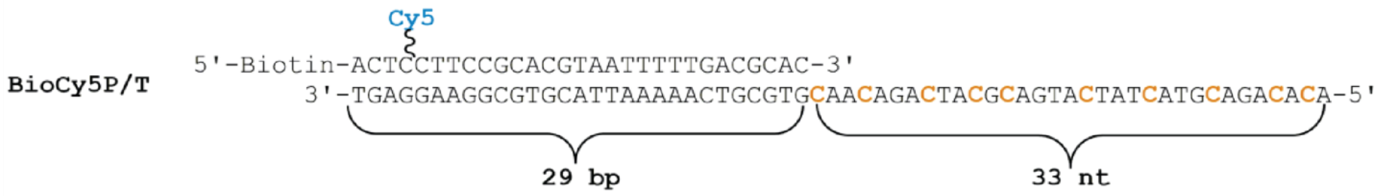


**b**

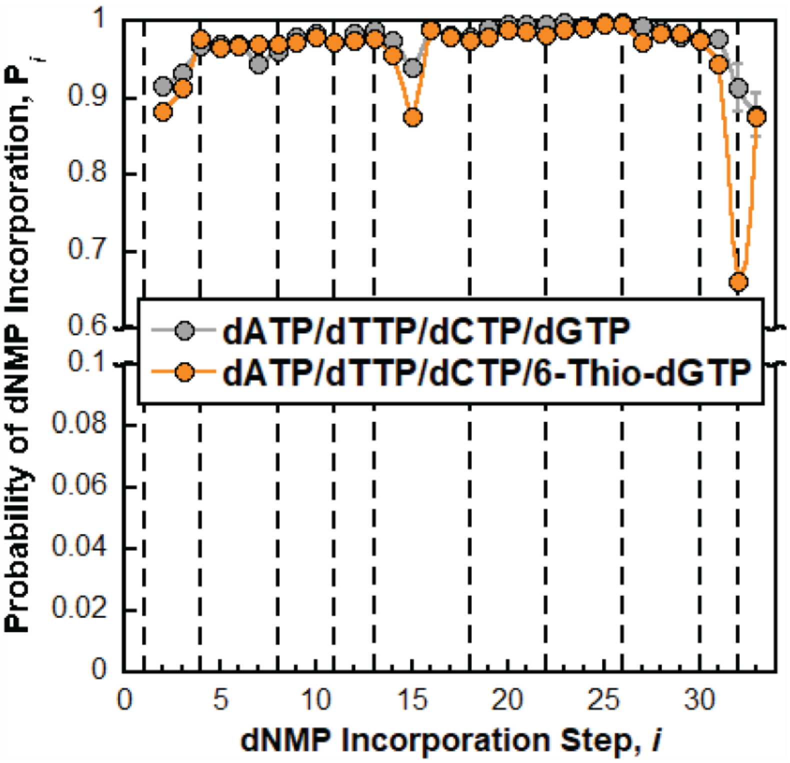

### Extended Figure 3

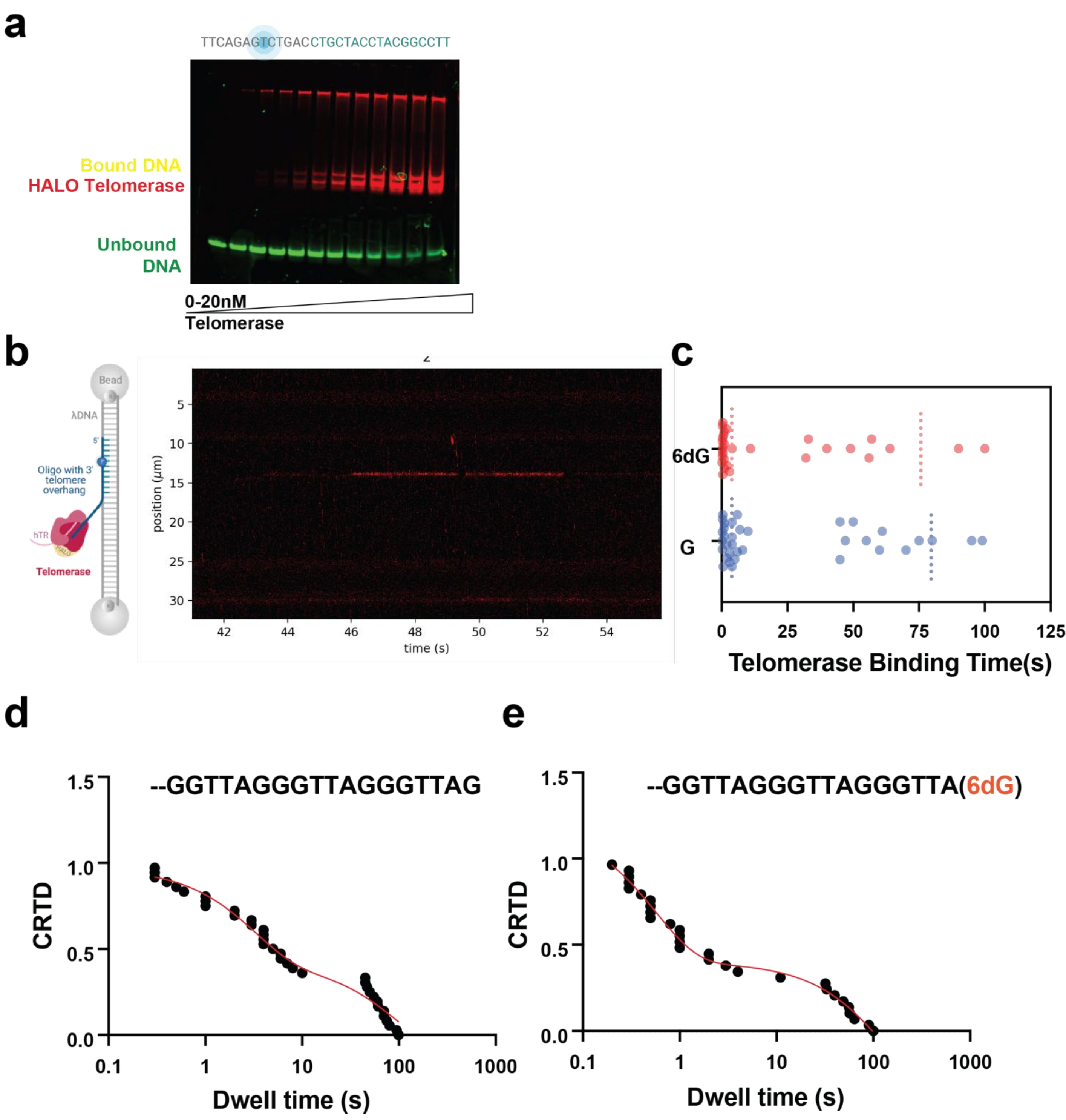

### Extended Figure 4

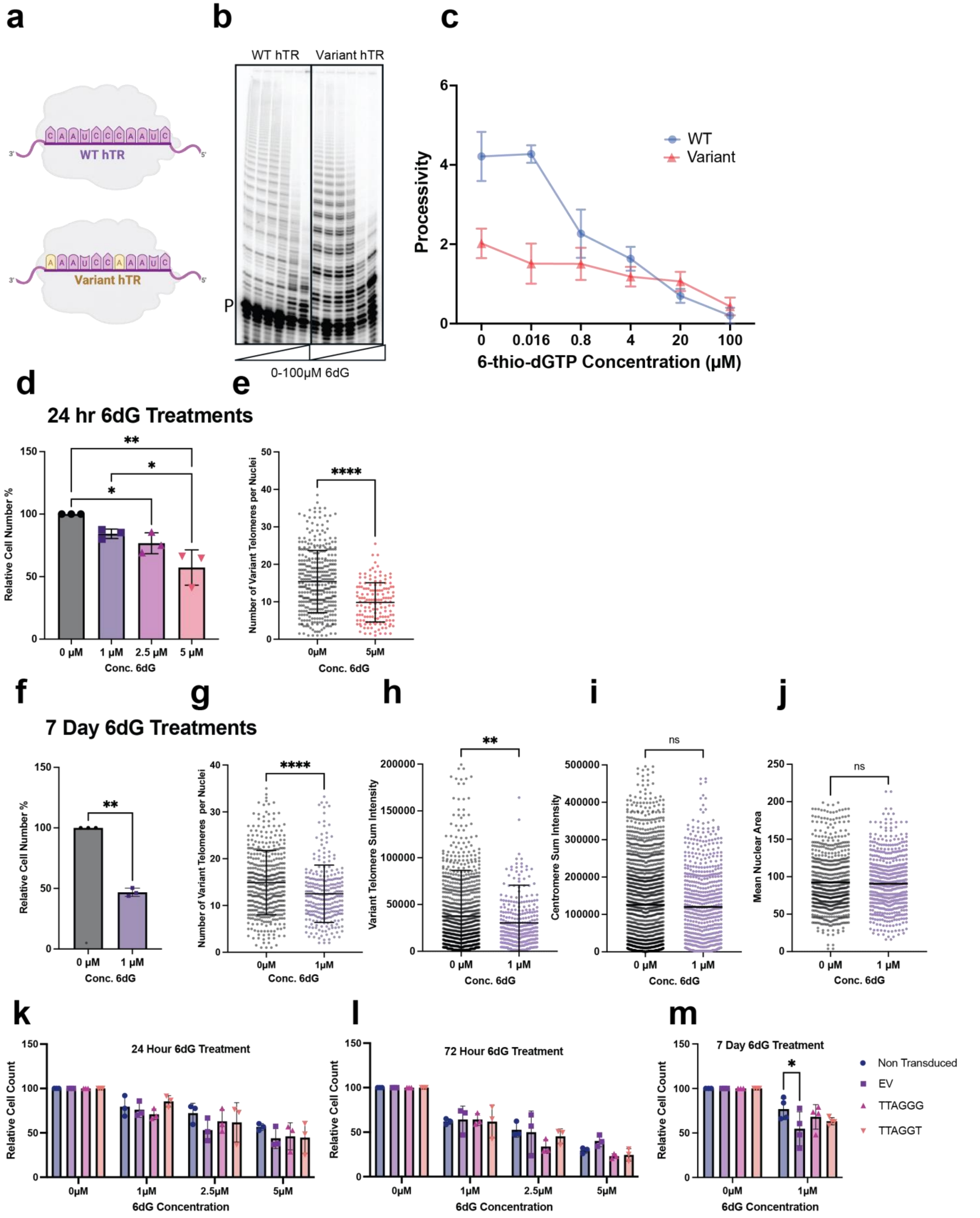

### Extended Figure 5

a

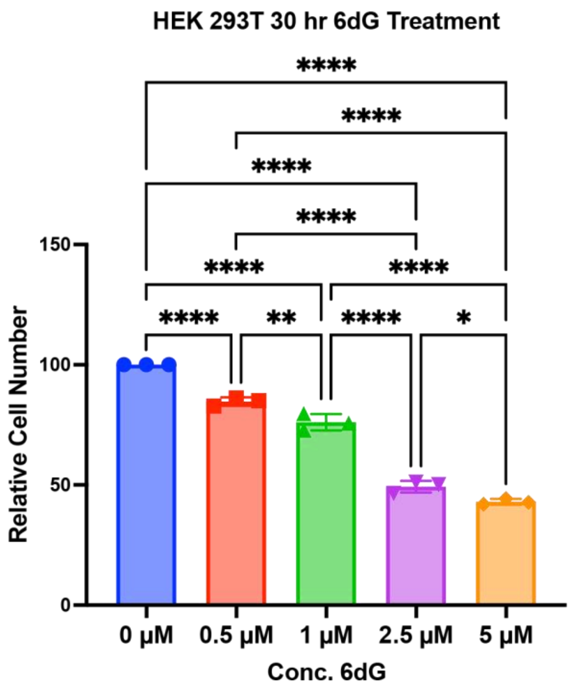


b

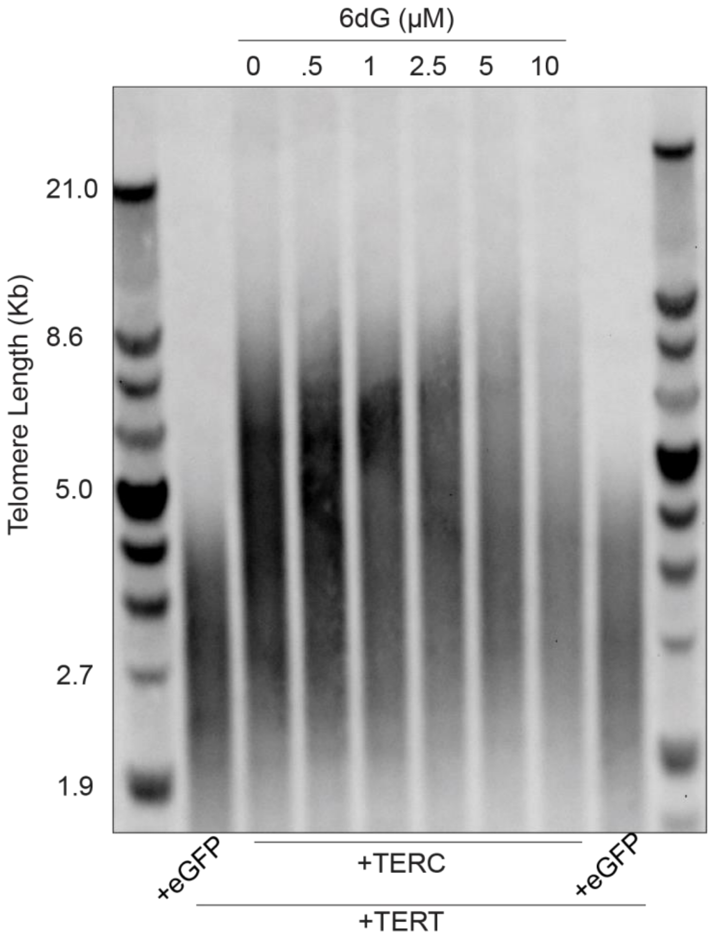


c

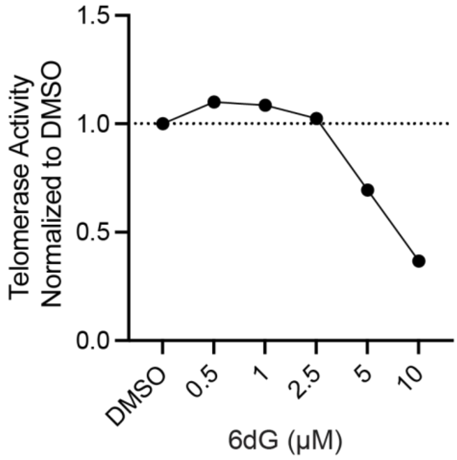

#### Extended Figure 6

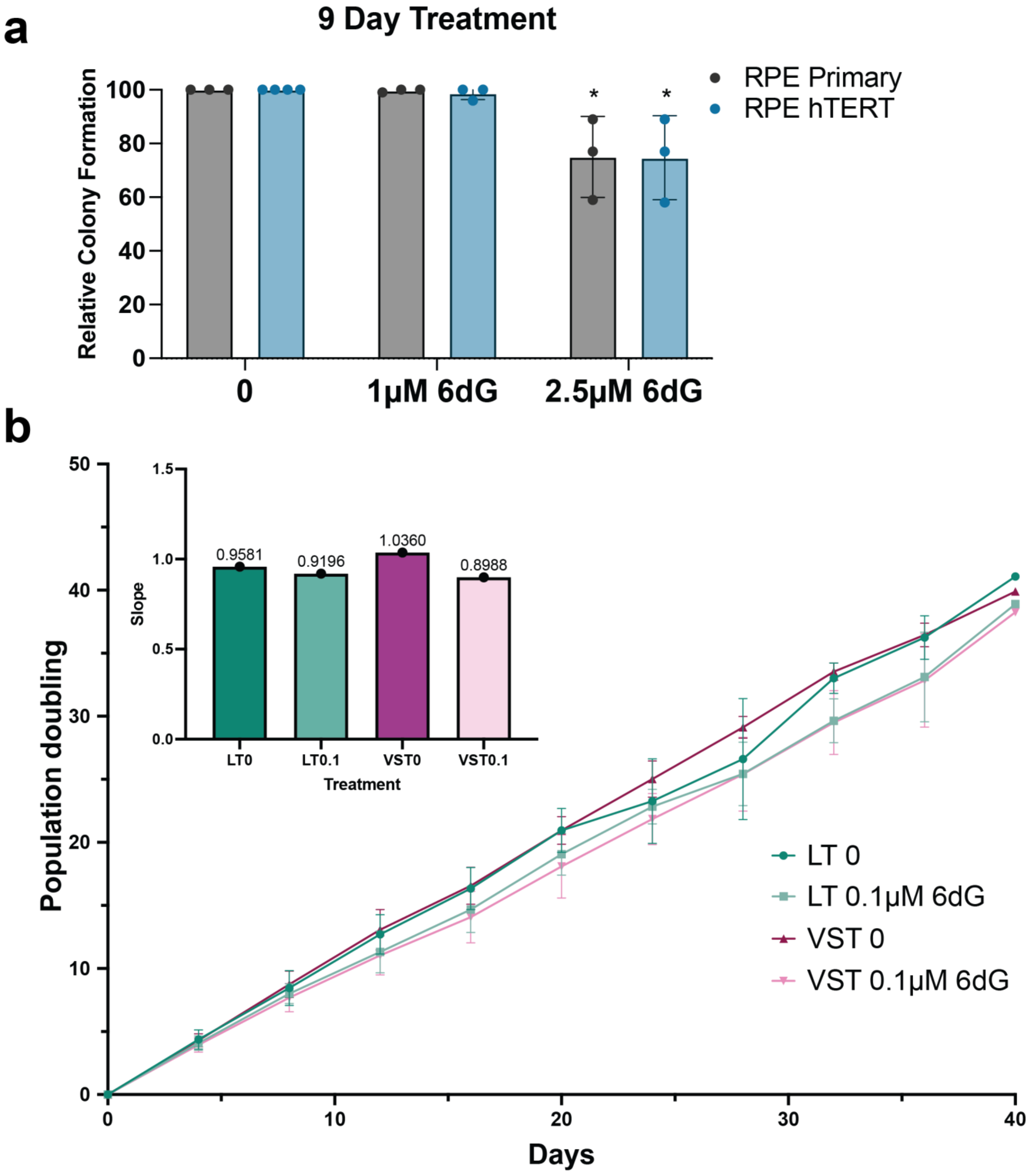

### Supplementary Table 1.

| Name | Sequence |
|---|---|
| **Oligo1** | GGTTAGGGTTAGGGTTAG |
| **Oligo1-6dGlast** | GGTTAGGGTTAGGGTTA/6dG/ |
| **Oligo1-6dGcentral** | GGTTAGGGTTAGG/6dG/TTAG |
| **Oligo2** | GTTAGGGTTAGGGTTAGG |
| **Oligo2-6dGlast** | GTTAGGGTTAGGGTTAG/6dG/ |
| **Oligo2-6dG2ndlast** | GTTAGGGTTAGGGTTA/6dG/G |
| **Oligo2-two6dGend** | GTTAGGGTTAGGGTTAG/6dG/6dG/ |
| **Primer A5** | TTAGGGTTAGCGTTAGGG |
| **EMSAoligo1** | TTCAGAG/iFluorT/CTGACGGTTAGGGTTAGGGTTAG |
| **EMSAoligo2** | TTCAGAG/iFluorT/CTGACGGTTAGGGTTAGGGTTA/6dG/ |
| **EMSAscramble** | TTCAGAG/iFluorT/ CTGACGGCTGCTACCTACGGCCTT |
| **hTR dot blot probe** | CGGTGGAAGGCGGCAGGCCGAGGC |
